# Haemosporidian infection does not alter aerobic performance in the Pink-sided Junco (*Junco hyemalis mearnsi*)

**DOI:** 10.1101/2021.09.20.460914

**Authors:** Maria Stager, Douglas K. Eddy, Zachary A. Cheviron, Matthew D. Carling

## Abstract

Avian haemosporidia are blood parasites that can have dramatic fitness consequences on their hosts, including largescale population declines when introduced to naïve hosts. Yet the physiological effects that accompany haemosporidian infection and underlie these fitness decrements are poorly characterized in most wild birds. Because haemosporidia destroy host red blood cells and consume host hemoglobin, they are predicted to have detrimental impacts on avian blood-oxygen transport and, as a result, reduce aerobic performance. However, the documented effects of infection on avian hematological traits vary across species and no effects have been demonstrated on avian aerobic performance to date. Here we quantified the physiological effects of haemosporidian infections on wild ‘Pink-sided’ Juncos (*Junco hyemalis mearnsi*) breeding in northwestern Wyoming, USA. We assayed hematological traits (hemoglobin concentration and hematocrit) and aerobic performance (resting and summit metabolic rates, thermogenic endurance, and aerobic scope), then screened individuals for haemosporidian infection *post-hoc* (*n* = 106 adult juncos). We found that infection status did not correlate with any of the physiological indices that we measured, suggesting there is little cost of haemosporidian infection on either junco aerobic performance or energy budgets. Our results highlight the need for more studies of haemosporidia infections in a broader range of species and in a wider array of environmental contexts.

## INTRODUCTION

Avian haemosporidian (the protozoan genera *Plasmodium, Haemoproteus*, and *Leucocytozoon*) are blood parasites, transmitted via insect vectors, that are responsible for causing avian malaria and other diseases. These obligate parasites are endemic across much of the globe and infect most bird families (Valkiunas 2005, LaPointe et al. 2012, Clark et al. 2014). The severity of haemosporidian infection can vary across avian hosts, with some species displaying resistance while others exhibit substantial fitness declines (Dawson and Bortolotti 2000, Merino et al. 2000, Marzal et al. 2005, Palinauskas et al. 2008, Kulma et al. 2013, Dadam et al. 2019). The novel introduction of the *Plasmodium*-carrying vector *Culex quinquefasciatus* to the Hawaiian Islands, for instance, devastated native bird communities (Scott et al. 1986). As a result, susceptible avifauna are now largely relegated to elevations in the archipelago above which the vector can persist (van Riper et al. 1986, Samuel et al. 2011).

Avian haemosporidia have been subject to investigation for more than a century but, due to advances in molecular techniques, the last twenty years have seen a surge in research focused on their prevalence, genetic diversity, ecology, and evolutionary biology (Clark et al. 2014, Rivero and Gandon 2018). Moreover, there has been renewed interest in characterizing the physiological effects of avian haemosporidia on their hosts. Much of this work has focused on the effect of infection on avian immune function (Ricklefs and Sheldon 2007, Bichet et al. 2012, Ellis et al. 2014, 2015), oxidative status (van de Crommenacker et al. 2012, Isaksson et al. 2013, Delhaye et al. 2016), telomere degradation (Asghar et al. 2015, Karell et al. 2017, Sudyka et al. 2019), stress hormones (Cornelius et al. 2014, Schoenle et al. 2018, 2019; Names et al. 2021), and feather growth patterns (Marzal et al. 2013, Coon et al. 2016). Far fewer studies have characterized the effects of haemosporidian infection on aerobic performance in wild birds.

Avian haemosporidia invade the red blood cells, as well as the organs and other tissues, of their hosts (Valkiunas 2005, Valkiūnas and Iezhova 2017). Because haemosporidia lyse erythrocytes and digest host hemoglobin as an amino acid source, they can have detrimental impacts on host blood-oxygen carrying capacity. Nonetheless, the documented effects of infection on hematological parameters, including hemoglobin concentration and hematocrit (the proportion of the blood volume comprised of red blood cells), vary across avian taxa. As expected, both hemoglobin concentration and hematocrit decrease with infection intensity (parasite load) among several species of Himalayan birds (Ishtiaq and Barve 2018). Similarly, Red-winged Blackbirds (*Agelaius phoeniceus*) exhibit increased hematological parameters with antimalarial treatment (Schoenle et al. 2017). However, no effect of haemosporidian infection was shown on hematocrit and/or hemoglobin concentration in experimentally-inoculated White-throated Sparrows (*Zonotrichia albicollis*; Kelly et al. 2020) or Great Reed Warblers (*Acrocephalus arundinaceus*; Hahn et al. 2018), captive Black-fronted Piping-guans (*Aburria jacutinga*; Motta et al. 2013), nor free-living American Redstarts (*Setophaga ruticilla*), Gray Catbirds (*Dumetella carolinensis*), Cedar Waxwings (*Bombycilla cedrorum*), or Red-eyed Vireos (*Vireo olivaceus*; Granthon and Williams 2017). Additionally, hematocrit was positively correlated with haemosporidian load in wild American Kestrels (*Falco sparverius*; Dawson and Bortolotti 1997). This variation highlights the potential importance of the underlying environmental context for evaluating these relationships and assessing the effects of infection.

Due to the vital importance of oxygen transport for aerobic metabolism, infection-induced reductions to blood-oxygen carrying capacity could negatively impact an individual’s aerobic performance (Hammond et al. 2000). Birds maintain a relatively high metabolic rate and frequently undertake aerobically taxing activities, such as active flight (Butler and Bishop 2000). Declines in hematocrit and hemoglobin due to haemosporidian infection are therefore predicted to have adverse effects on the ability of infected birds to perform even their most routine daily activities. In support of this hypothesis, captive canaries (*Serinus cararius*) inoculated with *P. relictum* exhibited reductions in both body temperature and metabolic rate in the cold (5°C) during the acute phase of infection, suggesting a decreased ability to thermoregulate – a key performance trait (Hayworth et al. 1987). If similar detriments to aerobic performance exist in infected birds in the wild, this could offer insight into how fitness effects manifest.

Even though haemosporidia may limit total aerobic capacity, parasite infections can simultaneously increase host energy budgets (e.g., Munger and Karasov 1994, Delahay et al. 1995, Careau et al. 2010, but see Robar et al. 2011). Individuals infected with haemosporidia could exhibit increased resting metabolic rates (an index of maintenance metabolic costs) due, for instance, to the costs of mounting an immune response or repairing damaged tissue (e.g., King and Swanson 2013). Increased energetic costs are associated with larger food requirements (Bozinovic and Sabat 2010), which may result in heightened exposure to predators, taxed food supplies, and reduced time for other fitness-enhancing activities. Accordingly, increased basal or resting metabolic rates are associated with reduced survival in some cases (Larivée et al. 2010, Nilsson and Nilsson 2016).

The range of metabolic rates available to an individual – a metric known as aerobic scope (Fry 1947) – may also be altered as a consequence of changes to either maximum aerobic capacity or resting metabolic rate. Generally a larger aerobic scope is thought to confer a greater ability to meet an energetic challenge or respond to environmental variation (Naya et al. 2012, Stager et al. 2016). Reductions to aerobic scope could result from either decreases to aerobic capacity (i.e., maximal or summit metabolic rates) and/or increases to resting or basal metabolic rates. Given that haemosporidian infection is predicted to influence both of these traits, avian aerobic scope could be greatly reduced in infected individuals.

To test these predictions, we quantified the physiological effects of haemosporidian infections on wild ‘Pink-sided’ Dark-eyed Juncos (*Junco hyemalis mearnsi*). Pink-sided Juncos breed in high-elevation forests of the Middle Rocky Mountains and spend the winter further south (Figure 1; Nolan, Jr. et al. 2020). Pink-sided Juncos may therefore come into contact with haemosporidia during the breeding season, as well as during the winter in the more southerly reaches of their distribution. Indeed, other Dark-eyed Junco populations are known to host haemosporidian infections (Slowinski et al. 2018, Becker et al. 2020). Across their distribution, Pink-sided Juncos are also subject to low temperatures throughout the year. Juncos and other small birds primarily rely on shivering thermogenesis to maintain a relatively constant body temperature when ambient temperatures drop below the thermoneutral zone, placing a premium on their aerobic performance in the cold (Swanson 1990, 2010). Juncos are therefore an appropriate system with which to explore the effects of haemosporidian infections on aerobic performance.

**Figure 1.**
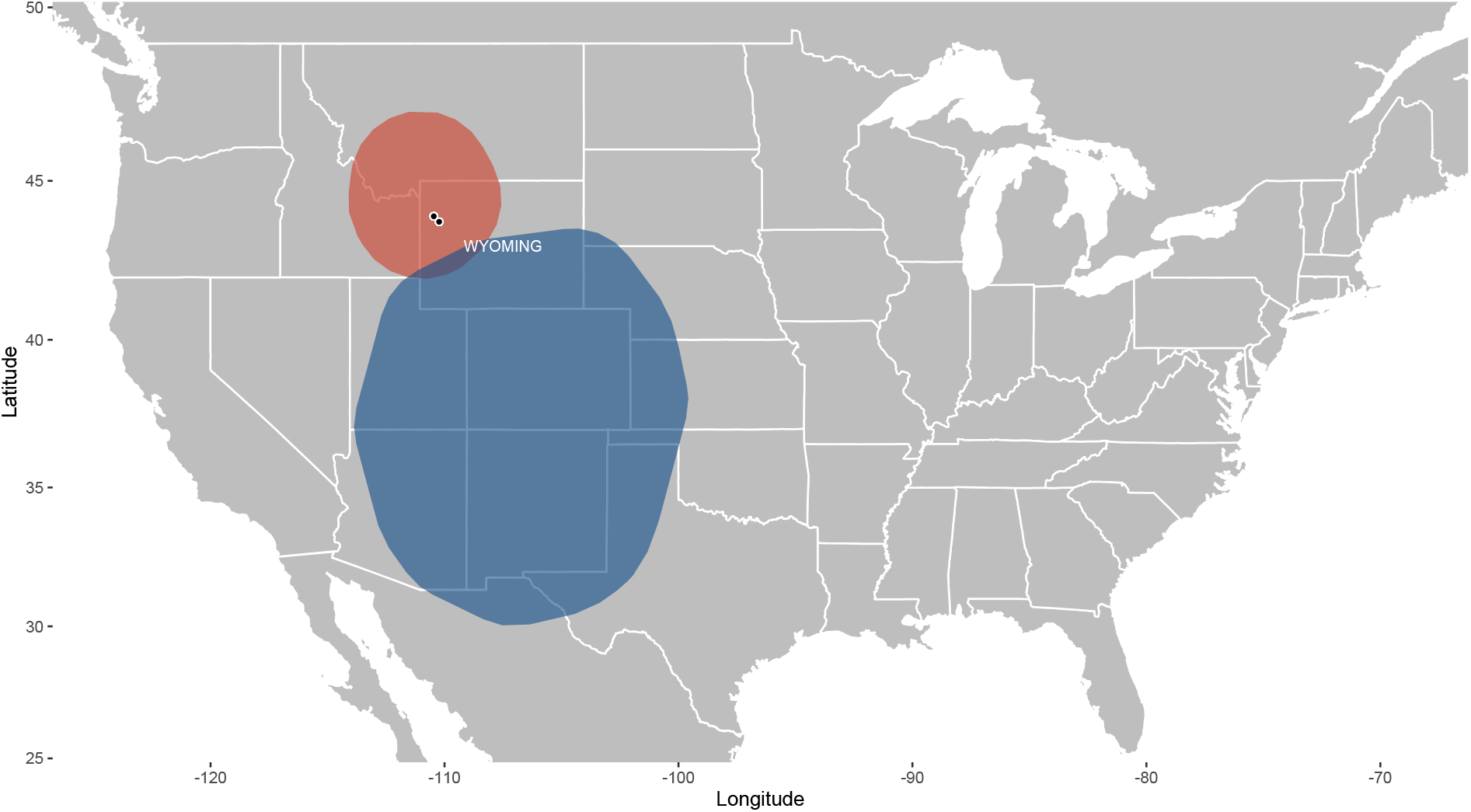
Approximate distribution of the Pink-sided Junco (*Junco hyemalis mearnsi*). Black dots denote our two sampling sites in northwest Wyoming. Shaded regions depict the 95% minimum convex polygons during the breeding (June-July; red) and non-breeding (December-February; blue) seasons. Minimum convex polygons were generated with the R package *adehabitatHR* (Calenge 2006) using *n* = 3243 breeding and *n* = 22614 non-breeding observations of Pink-sided Juncos made from 1947 to 2021 (eBird Basic Dataset 2021). The background map was plotted using R packages *ggplot2* (Wickham 2016), *mapdata* (Brownrigg 2018), and *maptools* (Bivand et al. 2021) with U.S. states outlined for reference. Polygons were added with package *ggforce* (Pederson 2021) using rounded corners (radius = 0.05).

We captured breeding Pink-sided Juncos at mid-elevation sites in northwestern Wyoming, USA. After bringing the juncos to a field laboratory, we measured hematocrit, hemoglobin concentration, summit metabolic rate (a measure of aerobic shivering capacity; Swanson 1990), thermogenic endurance in the cold, resting metabolic rate, and absolute aerobic scope. We assessed haemosporidian infection status of each individual *post hoc*. In line with the predictions outlined above, we expected that infected juncos would show decreased hematocrit, hemoglobin concentrations, summit metabolic rates, thermogenic endurance, and aerobic scope, and increased resting metabolic rates. With these data, our study provides an in-depth assessment of the effects of haemosporidian infections in wild, free-ranging birds.

## METHODS

### Field Capture

We sampled adult juncos on the breeding grounds at two sites in the Bridger-Teton National Forest in northwestern Wyoming, USA: Pacific Creek (PC; 43.92°N, 110.46°W; 2125 m.a.s.l.), and Togwotee Pass (TP; 43.75°N, 110.23°W; 2645 m.a.s.l). To attract birds, we played recordings of conspecific songs and calls and caught individuals in mist nets. We performed sampling in June over three years: we caught 20 individuals in 2013 (*n*_*PC*_ = 15; *n*_*TP*_ = 5), 36 individuals in 2015 (*n*_*PC*_ = 26; *n*_*TP*_ = 10), and 50 individuals in 2016 (*n*_*PC*_ = 29; *n*_*TP*_ = 21). We transported birds to laboratory facilities at the University of Wyoming / National Park Service Research Station in Grand Teton National Park within 2 h of capture. Birds were housed individually in cages (76.2 cm × 45.7 cm × 45.7 cm) and provided with food (meal worms and mixed seed) and water *ad libitum*. We held birds in captivity < 2 d (mean = 1.03 ±0.48), with the exception of 2 individuals in 2016 that were held for 3 d and have been removed from the physiological analyses. All animal procedures were performed with permission from University of Wyoming Institutional Animal Care and Use Committee (protocol #20150422MC00153-01).

### Metabolic Assays

For each individual, we quantified resting metabolic rate (the minimum oxygen consumption at rest, in the dark) and summit metabolic rate (the maximum oxygen consumption under cold exposure) using open-flow respirometry. Summit metabolic rate data have been previously published (Stager et al. 2021). Birds were not fasted before either measurement. At the start of each trial, individuals were weighed and then placed in a modified 1 L polycarbonate (Nalgene) chamber. An empty, identical chamber served as the baseline. Before each trial began, we allowed the respirometry system (described below) to run for > 1 h to equilibrate, then spanned the oxygen analyzer (FoxBox, Sable Systems International, Las Vegas, NV) at 20.95% O_2_ using baseline air.

Resting metabolic trials were conducted between 1900 and 0300 on the day of capture. Three individuals were run in a trial simultaneously and we assayed each individual for 1 h. All chambers were placed inside a temperature-controlled cabinet (PELT-5 temperature controller and PTC-1 temperature cabinet, Sable Systems International) held at 27°C (within the thermoneutral zone of summer-acclimatized juncos; Swanson 1991). Ambient air was scrubbed of water using Drierite (W. A. Hammond DRIERITE Co. LTD, Xenia, OH), pumped through a multi-channel mass flow meter (PP-2 Dual Channel Field Pump and Flowbar-8 Mass Flow Meter System, Sable Systems International), and directed into four chambers (animal or baseline) at 500 ml/min. We subsampled the excurrent air from each chamber, rotating among individuals at 20-min intervals and sampling the baseline chamber for 5-10 min every hour. We dried the excurrent air, scrubbed it of CO_2_ (Ascarite II, Thomas Scientific, Swedesboro, NJ), and dried it again before it entered the oxygen analyzer. Output was recorded using ExpeData software (ver. 0.2.57, Sable Systems International).

Summit metabolic trials were conducted the next day between 0730 and 1500 using a similar setup. For summit metabolic trials, we used heliox gas (21% oxygen/79% helium) equilibrated to atmospheric pressure and flow rates of 750 ml/min. Heliox facilitates heat loss at higher temperatures than is necessary in air, which helps avoid injury to experimental subjects (Rosenmann and Morrison 1974). We set the temperature cabinet to −5°C. A single individual was assayed at a time. In 2015, immediately following the trial, we verified hypothermia by inserting a thermocouple thermometer (Omega HH802U) into the individual’s cloaca. Body temperature was ≤36°C in all cases (compared to *μ* = 38.2 ±1.7°C following resting metabolic trials; *n* = 34). We did not take similar measures in 2013 or 2016; however, because summit metabolic rate can be been attained before the onset of hypothermia (Dutenhoffer and Swanson 1996), we did not exclude individuals from these other seasons in which hypothermia was not confirmed. Moreover, we did not find an effect of year on summit metabolic rate (see below).

We used custom R scripts to quantify metabolic traits from the resulting ExpeData files (https://github.com/Mstager/batch_processing_Expedata_files). We first used a linear correction to adjust for any drift that occurred in the baseline O_2_ value and calculated oxygen consumption (VO_2_; ml O_2_ per min) according to Eqn. 10.1 in Lighton (2008). We then calculated resting metabolic rate as the lowest mean VO_2_ averaged over a 10-minute period, and summit metabolic rate as the highest instantaneous VO_2_ averaged over a 5-min period. As a measure of thermogenic endurance, we also quantified the time that an individual maintained 90% or more of their summit metabolic rate (in min; Cheviron et al. 2013). We discarded measures characterized by large drift in baseline O_2_ (owing to ambient temperature fluctuations affecting the FoxBox) or inconsistent flow rates resulting in a total sample size of *n* = 81 individuals for resting metabolic rate and *n* = 93 for summit metabolic rate and thermogenic endurance. We also quantified absolute aerobic scope (summit metabolic rate – resting metabolic rate) for individuals for which we had both measures (*n* = 81).

### Blood and Tissue Sampling

Following summit metabolic trials, we collected a blood sample from the brachial vein to quantify hematological parameters. We quantified hematocrit in 2013 and 2015 by collecting ∼50 μl of blood in a microcapillary tube, centrifuging it for 5 min at 4,400 g, and measuring packed and total volumes with calipers. In 2015 and 2016, we quantified hemoglobin concentration (g/dL) by collecting 10 μl of whole blood in a cuvette and assaying it with a Hemocue Hb 201+ analyzer. In some cases, bleeds were insufficient to fill cuvettes/tubes or samples were destroyed in the centrifuge, resulting in a final sample size of *n* = 43 for hematocrit and *n* = 61 for hemoglobin. We then euthanized individuals via cervical dislocation, excised pectoralis tissue, and flash-froze it in liquid N_2_ for preservation. Individuals were later prepared as study skins and accessioned into the University of Wyoming Museum of Vertebrates (Table S1).

### Haemosporidia Detection

We detected haemosporidia infection by amplifying *Haemoproteus* and *Plasmodium* cytochrome *b* sequences from junco muscle tissue. We extracted DNA from pectoralis with a QIAcube^®^ robotic workstation (Qiagen, Valencia, Califormia) and DNeasy^®^ Blood & Tissue extraction kit (Qiagen, Valencia, California) using standard protocols. We then used primers HAEMNF (5’-CATATA TTAAGAGAA TTATGGAG-3’) and HAEMNR2 (5’-AGAGGTGTAGCATATCTATAC-3’) in anested PCR assay according to Waldenström et al. (2004). We performed all reactions in a 10-μl volume. For the first reaction, we used 6.45 μl dH_2_O, 1.0 μl (10 ng) genomic DNA, 1 μl MgCl_2_ (2.5 mM), 1.0 μl 10X PCR buffer, 0.25 μl dNTPs (10mM each dNTP), 0.1 μl of each primer, and 0.1 μl *Taq* (5,000 units/ml; New England BioLabs, Ipswitch, Massachusetts). We performed denaturation at 95°C for 3 min; 20 cycles of denaturation/elongation at 96°C for 30 sec, 49°C for 30 sec, and 75°C for 45 sec; followed by a final elongation step at 72°C for 10 min. For the second PCR in the nested assay, we used the same quantities of reagents and 1.0 μl of PCR product from the previous reaction. We performed denaturation at 95°C for 3 min; 35 cycles of denaturation/elongation at 96°C for 30 sec, 49°C for 30 sec, and 75°C for 45 sec; followed by a final elongation step at 72°C for 10 min. This assay amplified a 524 bp region of *Haemoproteus* and *Plasmodium* cytochrome *b*. We ran the final PCR product on a 1% agarose gel. We ran positive and negative controls for all PCRs, and we performed three reactions for each individual in order to detect weak infections and to account for low levels of parasite DNA in the sample.

### Statistical Analyses

All analyses were performed in the R statistical environment (ver. 4.0.2, R Core Team 2018). We quantified relationships among physiological traits to determine whether associations between hematological parameters and aerobic performance existed. Metabolic traits are known to correlate with body mass, so we first quantified this relationship using a linear regression and included body mass as a covariate in all subsequent analyses. Because we collected more measures of hemoglobin concentration than hematocrit and the two metrics were correlated within individuals (see Results below), we chose to assess the correlation between hemoglobin concentration and metabolic traits but not that of hematocrit and metabolic traits. We did so using linear regressions with each metabolic trait as the response variable, and hemoglobin concentration and body mass as covariates.

Because our sampling scheme spanned three years and two sites, we also tested for differences among sites and years for each of the physiological traits. We performed Welch’s two sample t-tests to test for differences among sites or years for hemoglobin concentration and hematocrit. We used linear regressions to test the effect of site or year on each of resting metabolic rate, summit metabolic rate, aerobic scope, and thermogenic endurance, with body mass again as a covariate.

We used generalized linear models with a binomial response to test for differences in infection status (infected vs. not infected) among sites and years. We performed Welch’s two sample t-tests to test for differences among infection statuses for each hemoglobin concentration and hematocrit. To test the effect of haemosporidian infection on metabolic traits (resting metabolic rate, summit metabolic rate, aerobic scope, thermogenic endurance), we composed linear regressions with each trait as the response variable and infection status as a predictor, as well as body mass, year, and site as covariates where appropriate.

## RESULTS

### Relationships among physiological traits

Body mass positively correlated with resting metabolic rate (*β* = 0.05 ± 0.01 ml O_2_ g^-1^ min^-1^, *p* = 2.93 × 10^−5^), summit metabolic rate (*β* = 0.25 ± 0.08 ml O_2_ g^-1^ min^-1^, *p* = 1.6 × 10^−3^), thermogenic endurance (*β* = 3.42 ± 1.08 min, *p* = 2.02 × 10^−3^), and aerobic scope (*β* = 0.22 ± 0.09 ml O_2_ g^-1^ min^- 1^, *p* = 0.01) and was therefore included as a covariate in subsequent analyses (Figure 2). Importantly, hemoglobin concentration and hematocrit positively correlated (*r* = 0.69) in the 32 individuals for which we collected both measures in 2015. Higher hemoglobin concentrations were also associated with higher summit metabolic rate and aerobic scope, and lower resting metabolic rate (Table 1).

**Figure 2.**
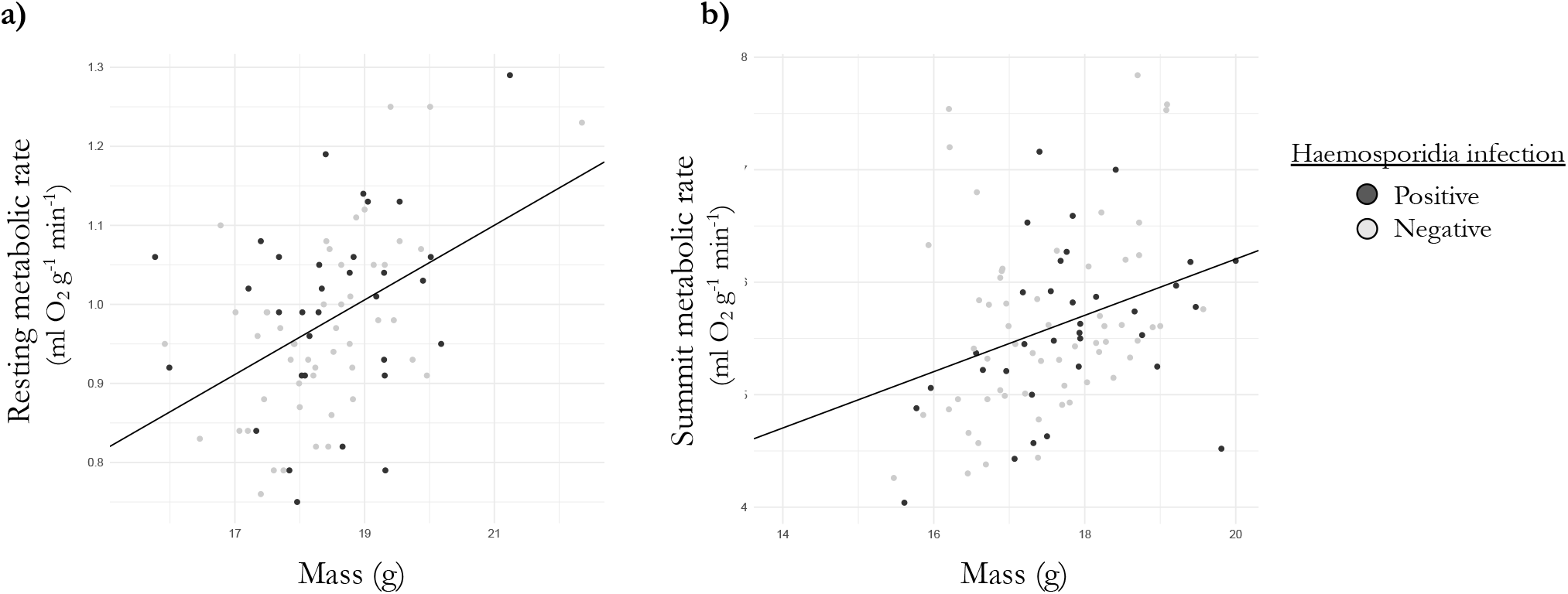
Metabolic traits are positively correlated with body mass but are not affected by haemosporidia infection. a) Resting metabolic rate. b) Summit metabolic rate. Dots represent individuals, color-coded by haemosporidia infection status: positive (dark grey) or negative (light grey).

**Table 1.** Effects of hemoglobin concentration on metabolic traits resulting from a linear regression with body mass as a covariate.

| Trait | df | Hemoglobin |  |  | Body Mass |  |  | Adj. R<sup>2</sup> |
| --- | --- | --- | --- | --- | --- | --- | --- | --- |
| | | $\beta$ | SE | $p$ | $\beta$ | SE | $p$ | |
| Resting metabolic rate | 53 | -0.02 | 0.004 | $1.56 \times 10^{-3}$ | 0.07 | 0.01 | $2.85 \times 10^{-7}$ | 0.42 |
| Summit metabolic rate | 58 | 0.07 | 0.03 | 0.04 | 0.29 | 0.09 | $1.37 \times 10^{-3}$ | 0.20 |
| Thermogenic endurance | 58 | -0.05 | 0.59 | 0.93 | 4.67 | 1.65 | $6.48 \times 10^{-3}$ | 0.09 |
| Aerobic scope | 53 | 0.07 | 0.03 | 0.04 | 0.31 | 0.08 | $6.35 \times 10^{-4}$ | 0.26 |

### Physiological variation across sites and years

Hemoglobin concentration did not differ among sites (*t* = 0.26, df = 43, *p* = 0.80) but it did differ between the two years that we collected it (*t* = −4.32, df = 58, *p* = 6.18 × 10^−5^): individuals sampled in 2016 had higher hemoglobin concentrations (*n* = 28, *μ*= 18.25 g/dL) than those sampled in 2015 (*n* = 32, *μ*= 15.87). Conversely, hematocrit differed across sites (*t* = 2.71, df = 21, *p* = 0.01), with individuals at PC exhibiting higher hematocrit (*n* = 30, *μ*= 0.51) than those at TP (*n* = 13, *μ*= 0.46), but it did not differ between the two years (2013 and 2015) that we collected it (*t* = −0.91, df = 18, *p* = 0.38). Resting metabolic rate also differed across years (*β*_2015_ = −0.07 ± 0.04, *p* = 0.12; *β*_2016_ = −0.12 ± 0.04, *p* = 3.44 × 10^−3^) but summit metabolic rate, thermogenic endurance, and aerobic scope did not (Table 2). Metabolic traits did not differ across sites (Table 3).

**Table 2.** Effects of Year on metabolic traits from a linear regression with body mass as a covariate. 2013 served as the reference year.

| Trait | df | Year<sub>2015</sub> |  |  | Year<sub>2016</sub> |  |  | Body Mass |  |  | Adj. R<sup>2</sup> |
| --- | --- | --- | --- | --- | --- | --- | --- | --- | --- | --- | --- |
| | | $\beta$ | SE | $p$ | $\beta$ | SE | $p$ | $\beta$ | SE | $p$ | |
| Resting metabolic rate | 77 | -0.07 | 0.04 | 0.12 | -0.12 | 0.04 | $3.44 \times 10^{-3}$ | 0.04 | 0.01 | $1.99 \times 10^{-4}$ | 0.28 |
| Summit metabolic rate | 89 | -0.15 | 0.23 | 0.51 | -0.11 | 0.23 | 0.64 | 0.25 | 0.08 | $3.33 \times 10^{-3}$ | 0.08 |
| Thermogenic endurance | 89 | 3.02 | 3.06 | 0.33 | -4.34 | 3.03 | 0.16 | 2.45 | 1.09 | 0.03 | 0.16 |
| Aerobic scope | 74 | -0.31 | 0.31 | 0.32 | -0.19 | 0.31 | 0.55 | 0.23 | 0.09 | 0.01 | 0.06 |

**Table 3.**
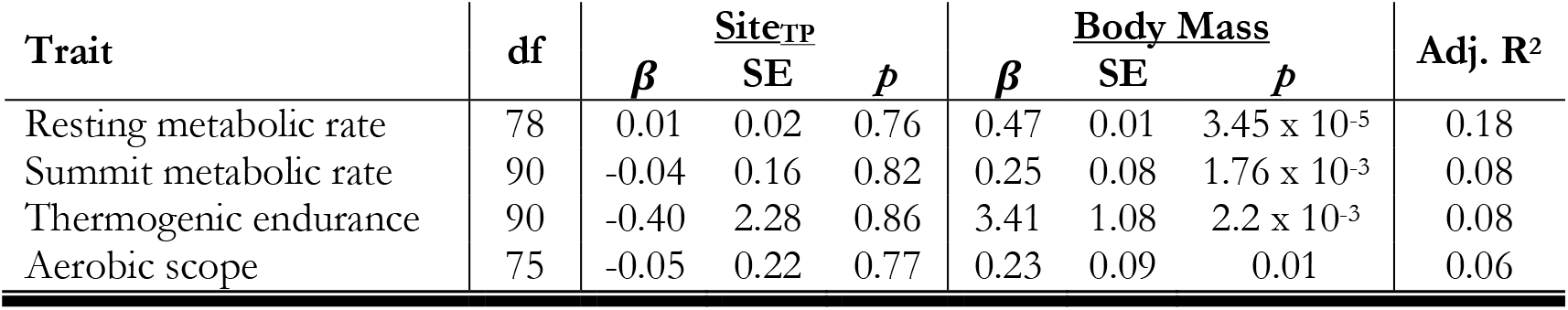
Effects of Site on metabolic traits resulting from a linear regression with body mass as a covariate. PC served as the reference site.

| Trait | df | Site<sup>TP</sup> |  |  | Body Mass |  |  | Adj. R<sup>2</sup> |
| --- | --- | --- | --- | --- | --- | --- | --- | --- |
| | | $\beta$ | SE | *p* | $\beta$ | SE | *p* | |
| Resting metabolic rate | 78 | 0.01 | 0.02 | 0.76 | 0.47 | 0.01 | 3.45 x 10<sup>-5</sup> | 0.18 |
| Summit metabolic rate | 90 | -0.04 | 0.16 | 0.82 | 0.25 | 0.08 | 1.76 x 10<sup>-3</sup> | 0.08 |
| Thermogenic endurance | 90 | -0.40 | 2.28 | 0.86 | 3.41 | 1.08 | 2.2 x 10<sup>-3</sup> | 0.08 |
| Aerobic scope | 75 | -0.05 | 0.22 | 0.77 | 0.23 | 0.09 | 0.01 | 0.06 |

### Effects of haemosporidian infection on physiological traits

We detected haemosporidian infection in 41 of 106 juncos that we sampled. Infection status did not differ across sampling sites (*β* = −0.35 ± 0.43, *p* = 0.42) but it did differ across years, with infection rate being highest in 2015 (*β*_2013_ = −1.07 ± 0.59, *p* = 0.07; *β*_2016_ = −1.07 ± 0.46, *p* = 0.02; Figure 3). Haemosporidian infection status, however, did not influence hemoglobin concentration in 2015 (*t* = −1.41, df = 28, *p* = 0.17), 2016 (*t* = −0.14, df = 8, *p* = 0.89), or when both years were combined (*t* = −0.01, df = 54, *p* = 0.99). Similarly, infection status did not influence hematocrit within either site (PC: *t* = −0.07, df = 24, *p* = 0.95; TP: *t* = −0.41, df = 4, *p* = 0.70) or when both sites were combined (*t* = −0.27, df = 41, *p* = 0.79). Nor did infection status correlate with resting metabolic rate (*β* = 0.01 ± 0.02, *p* = 0.67), summit metabolic rate (*β* = −0.11 ± 0.16, *p* = 0.48), thermogenic endurance (*β* = − 0.79 ± 2.28, *p* = 0.73), or aerobic scope (*β* = −0.10 ± 0.17, *p* = 0.56).

**Figure 3.**
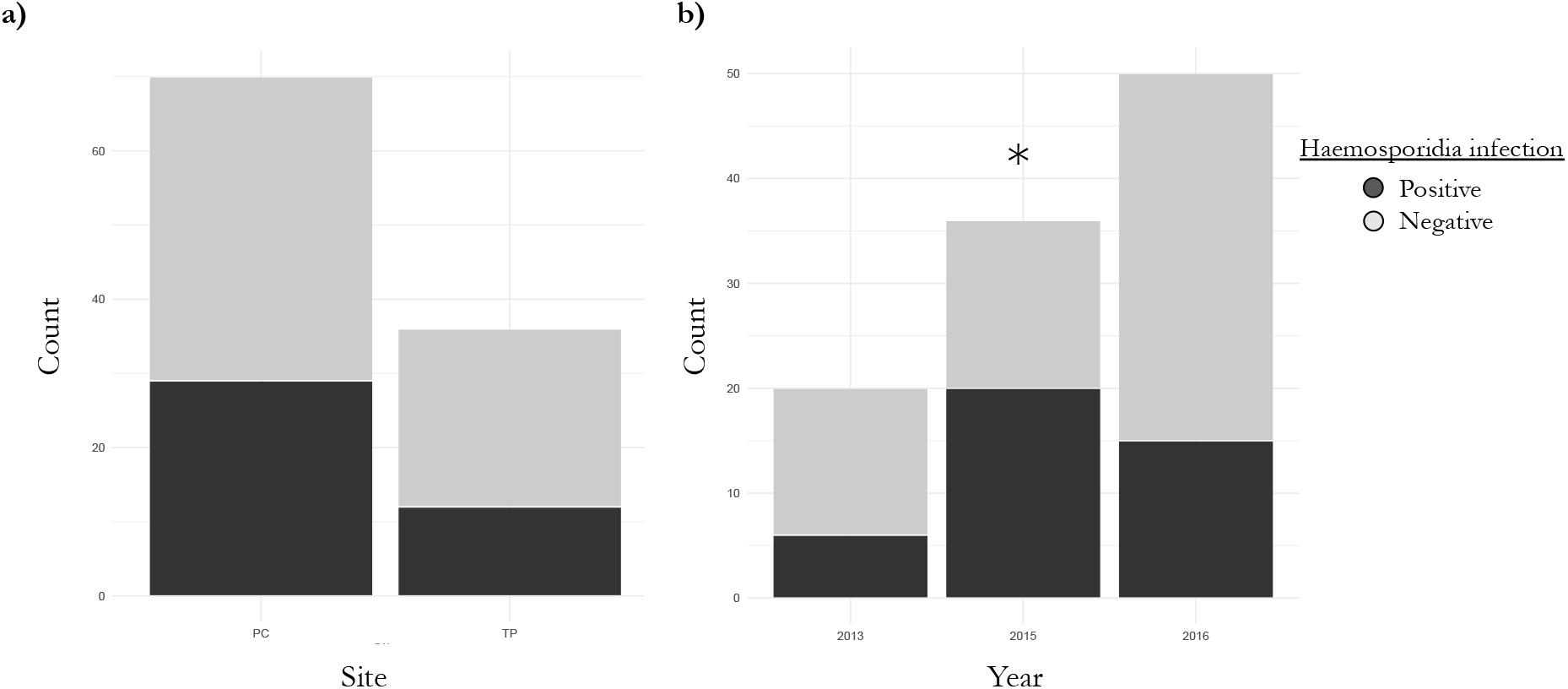
Haemosporidian infection counts across a) sampling sites and b) years. Colors indicate positive (dark grey) or negative (light grey) infection status. Asterisk denotes statistical significance (p < 0.05) from generalized linear model.

## DISCUSSION

Haemosporidian infections can negatively impact avian survival but the physiological mechanisms that contribute to these fitness consequences have only recently begun to be explored in free-living birds. We found that haemosporidian infections were prevalent in a wild population of adult Pink-sided Juncos at a rate of 39% during the breeding season. However, we failed to find correlations between infection status and junco hematological parameters (hemoglobin concentration and hematocrit) or metabolic traits (resting metabolic rate, summit metabolic rate, absolute aerobic scope, or thermogenic endurance). This work suggests there is little cost of haemosporidian infection on either junco aerobic performance or energy budgets.

Haemosporidia can reduce host red blood cell numbers through direct hemolysis (Valkiunas 2005). Accordingly, treating birds with antimalarials (Schoenle et al. 2017) or applying pesticides (to reduce the presence of haemosporidian vectors; Krams et al. 2013) has resulted in increased hemoglobin concentrations in treated individuals. We found, however, that haemosporidian infection was not associated with reductions in either of the junco red blood cell parameters that we measured. This finding is consistent with several other studies that did not demonstrate an association between infection status (infected vs. not infected) and hematocrit and/or hemoglobin (Motta et al. 2013, Granthon and Williams 2017, Hahn et al. 2018, Kelly et al. 2020). Our sampling occurred at mid elevations, but it is possible that potential negative effects of infection on blood parameters are more likely to materialize at higher elevations (e.g., Ishtiaq and Barve 2018) where the low partial pressure of oxygen may place a greater premium on blood O2 carrying capacity to support aerobic performance. Additionally, previous exposure to haemosporidia can shape hematological responses during secondary infection. Experimentally-inoculated Canaries, for example, exhibited reduced hematocrit during primary infection but not secondary infection (Cellier-Holzem et al. 2010). These examples highlight the potential complicated nature of the relationships between hematological parameters and haemosporidian infection.

One possible consequence of reductions to blood-oxygen transport is limited aerobic performance (Hammond et al. 2000). Indeed, we found that junco hemoglobin concentration was positively correlated with both summit metabolic rate and thermogenic endurance. Both of these metrics contribute to an individual’s resistance to cold (Swanson 2001) and are likely to positively correlate with survival under low ambient temperatures (Petit et al. 2017). Minimum temperatures at our sampling sites averaged 4°C during our capture period (range: −2.5 – 6.5°C; Thornton et al. 2016), indicating that cold tolerance may be important for survival even during the breeding season for Pink-sided Juncos. However, because we did not observe a negative effect of haemosporidia on hematological parameters, it is perhaps unsurprising that we also failed to find an association between infection status and either measure of aerobic performance. Similarly, a recent pair of studies using Great Reed Warblers did not find any indication that either acute or chronic haemosporidian infections substantially influenced an individual’s maximum metabolic rate, endurance, or migratory performance (Hahn et al. 2018, Emmenegger et al. 2021). Thus, in all three studies that we know of using wild birds (including our own), empirical support is lacking for predicted connections between haemosporidian infection and avian aerobic performance.

An alternative route by which parasites are predicted to reduce host fitness is through increases to host energy budget (reviewed in Robar et al. 2011). For example, host oxidative stress resulting from haemosporidian infection (e.g., van de Crommenacker et al. 2012, Isaksson et al. 2013) is hypothesized to be mediated through increased energy requirements (Delhaye et al. 2016). The specific effects of haemosporidian parasites on avian energetics have been little explored; however, the few studies that exist do not support the prediction that host energy budgets increase with infection. For instance, resting metabolic rate decreased slightly during the crisis period of *P. relictum* infection in canaries (Hayworth et al. 1987). Moreover, haemosporidian infection did not affect resting metabolic rates of Great Reed Warblers (Hahn et al. 2018). We similarly found that junco resting metabolic rate was not influenced by infection status. Taken together, this suggests that any energetic costs associated with mounting avian immune defenses in response to haemosporidian infection are not great enough to be reflected in resting metabolic rates.

In the absence of changes to either summit or resting metabolic rates, we accordingly did not observe an effect of infection on aerobic scope. We are not aware of any other studies that have investigated relationships between aerobic scope and either parasite infection (of any kind) in birds or haemosporidian infection in non-avian hosts. However, other parasitic infections are associated with declines in aerobic scope in several non-avian systems (Kumaraguru et al. 1995, Careau et al. 2012, Bruneaux et al. 2017, Hvas et al. 2017, Wilde et al. 2019). For example, Botfly (*Cuterebridae*) larval infection of deer mice (*Peromyscus maniculatus*) reduced host aerobic scope (via decreased summit metabolic rate) and was associated with lower host daily survival rates (Wilde et al. 2019). More studies characterizing the effects of parasites on aerobic scope are thus needed to test the generality of our findings in other birds.

One potential shortcoming of our work is that we did not quantify parasitemia and, thus, cannot determine whether haemosporidian intensity affects junco physiology. Furthermore, because we measured wild birds with naturally occurring infections, it is unclear where or when they acquired haemosporidia. Although we can personally attest that the breeding grounds were full of biting insects, individuals of this subspecies overwinter at latitudes where insect vectors are also likely to occur and thus exposure could transpire year-round. The time course of avian haemosporidia infection begins with a brief period of high parasitemia (the acute phase), followed by lower parasitemia that can last weeks, months or years (the chronic phase) during which the host is subject to relapse (Asghar et al. 2012). Physiological effects should be strongest during the acute phase of infection, but due to its short duration, naturally infected individuals are less likely to be sampled during this time. Notably, Hahn et al. (2018) did not document physiological effects of infection during either the acute or chronic stage in either captive or free-living birds. However, we cannot discount the fact that, if birds in this study exhibited chronic infections, juncos may still show pronounced physiological effects during the acute stage of infection. Another possibility is that some proportion of the population did suffer from impaired performance and died as a result. Severely affected individuals are difficult to sample (Valkiunas 2005), and so our dataset is likely biased to only those individuals that survived initial stages of infection and were capable of responding to our playback. Thus, we may have excluded individuals that did experience stronger physiological effects. Experimental inoculations can be useful in this regard (Kelly et al. 2020), but even some recent experiments have failed to find effects of haemosporidian infection (Hahn et al. 2018).

Ultimately, the effects of infection may differ between species with long coevolutionary histories and those that have come into contact more recently. In contrast to avian populations that have witnessed declines following the novel introduction of haemosporidian vectors, like those of Hawaii, the effects of infection on host fitness are often more subtle in populations that have coevolved with haemosporidia for a long time (LaPointe et al. 2012). What is more, a broad scale study of the relationship between haemosporidia diversity and avian diversity in the tropical Andes of South America suggests that haemosporidia do not influence avian species turnover, calling into question their effects on species’ competitive capabilities and distributions (McNew et al. 2021). In this context, it may not be surprising that we show that haemosporidian infection does not influence aerobic performance or energetics in a species for which haemosporidia are likely endemic. Our findings therefore point to the need for more detailed studies of the effects of haemosporidia infections on a broader range of species in a wider array of environmental contexts.

## ACKNOWLEDGEMENTS

Thank you to Chelsea Maguire for helping to perform PCR and to Nathan Senner, Rachel Fanelli, Brittany Nordberg, Ashleigh Rhea, and Zac Swope for helping in the field. This work was funded by grants from the UW/NPS Research Station to MDC and ZAC, and from the Wilson Ornithological Society to DKE. Further field work support was provided by the University of Wyoming Museum of Vertebrates.

## Supplemental Materials

Individual-level data including capture date, sampling site, and accession number are included in .csv format in Table S1.

## Notes

### Competing Interest Statement

The authors have declared no competing interest.

